# DAB-APT: a Fluorescence-Based Assay for Determining Aminopropyl Transferase Activity and Inhibition

**DOI:** 10.1101/2024.04.09.588734

**Authors:** Pallavi Singh, Jae-Yeon Choi, Choukri Ben Mamoun

**Affiliations:** Department of Internal Medicine, Section of Infectious Diseases, Yale School of Medicine, New Haven, CT, USA

**Author notes:** Correspondence: Choukri Ben Mamoun.

## Abstract

Polyamines are polycationic molecules that are crucial in a wide array of cellular functions. Their biosynthesis is mediated by aminopropyl transferases (APTs), promising targets in antimicrobial, antineoplastic and antineurodegenerative therapies. A major limitation, however, is the lack of high-throughput assays to measure their activity. We developed the first fluorescence-based assay, DAB-APT, for measurement of APT activity using 1,2-diacetyl benzene, which forms fluorescent conjugates with putrescine, spermidine and spermine with fluorescence intensity increasing with increasing carbon chain length. The assay has been validated using APT enzymes from *S. cerevisiae* and *P. falciparum* and is suitable for high-throughput screening of large chemical libraries. Given the importance of APTs in infectious diseases, cancer and neurobiology, our DAB-APT assay has broad applications, holding promise for advancing research and drug discovery efforts.

## Main

Polyamines are polycationic aliphatic biogenic molecules containing carbon chains of varying lengths and different number of amino groups. They are found ubiquitously in all eukaryotic and prokaryotic organisms and are essential for cell growth, differentiation, and survival ^1,2^. The most common polyamines include putrescine (diamine), spermidine (triamine), and spermine (tetramine)^2^. The pathway for the biosynthesis of polyamines has long been considered an attractive target for the development of novel therapies for the treatment of microbial infections, cancer, and neurodegeneration ^3–7^. In various organisms, including *Plasmodium falciparum*, the main causative agent of human malaria, the polyamine biosynthesis pathway initiates with the decarboxylation of ornithine via ornithine decarboxylase (ODC) to form putrescine ^8,9^. Spermidine synthase (SPDS), a member of the aminoproyltransferase (APT) class of enzymes, then catalyzes the transfer of the propylamine group from decarboxylated S-adenosyl methionine (dc-SAM) to putrescine to form spermidine^10^. Spermidine can also accept an aminopropyl group derived from dc-SAM to form spermine, a reaction catalyzed by a second APT enzyme, spermine synthase (SPMS). In yeast, SPDS and SPMS activities are catalyzed by the Spe3 and Spe4 enzymes. Genetic studies demonstrated that disruption of *SPE3* gene results in spermidine, spermine, β-alanine, or pantothenic acid auxotrophy, whereas loss of *SPE4* results in spermine, β-alanine, or pantothenic auxotrophy ^11^. These findings highlight the importance of polyamine biosynthesis as an attractive target for the development of new antimicrobial drugs. Unlike *P. falciparum* and *S. cerevisiae*, other lower eukaryotes, such as *Babesia* and *Eimeria* species, lack an ODC enzyme and rely exclusively on the uptake of polyamines for survival^12,13^. While polyamines have been implicated in various cellular functions, data available so far suggest that one of the crucial functions of the polyamine biosynthesis pathway is to form spermidine, which serves as a precursor for the synthesis of hypusine, an uncommon but critical amino acid for the activity of the eukaryotic translation factor eIF5A^14^.

Although APT enzymes involved in polyamine biosynthesis have been known for many years to be attractive targets for the development of new antimicrobials, means to inhibit their activity have relied primarily on the use of substrate analogs such as α-difluoromethylornithine (DFMO; ornithine analog) and 1-aminooxy-3-aminopropane^15,16^. A search for new small molecules or natural products with specific and more potent activity has been hampered by the lack of assays that are amenable to high-throughput screening of chemical libraries. Measurement of the activity and catalytic parameters of APT enzymes has so far relied mainly on the use of radioisotope-based assays, involving radiolabeled substrates, such as radiolabeled [^14^C]Spermine trihydrochloride and decarboxylated S-adenosyl[methyl-^3^H]methionine, which can be incorporated into the target substrate by the aminopropyl transferase^17,18^. A second method involved dansylation of polyamines synthesized through reactions with dansyl chloride, followed by quantitative analysis using high-performance liquid chromatography (HPLC)^19^. A third approach involved capillary electrophoresis with laser-induced fluorescence (CE-LIF) by derivatizing spermidine with 7-fluoro-4-nitrobenzo-2-oxa-1,3-diazole (NBD-F)^20^. Finally, spermidine synthase activity was assessed using a specific monoclonal antibody against the reaction product, 5’-methylthioadenosine (MTA), coupled with a homogeneous time-resolved fluorescence technique^21^. While these four assays are sensitive, they are low-throughput and involve time-consuming procedures. Together these data highlight the need for a new assay that could be used in a high throughput format to screen chemical libraries.

Here, we report the development of a simple and easy to use assay (DAB-APT) for the measurement of APT enzyme activity. The assay, which can be used in 96- and 384-well formats uses the primary amine reactive 1,2-diacetyl benzene, which forms fluorescent conjugates with putrescine, spermidine and spermine with fluorescence intensity increasing with the length of each of these polyamines. The DAB-APT assay was evaluated and validated using the Spe3 and Spe4 enzymes of *S. cerevisiae*, and the SPDS enzyme of *P. falciparum* and shown to be suitable for determining their biochemical activity, catalytic properties, and inhibition by known inhibitors, setting the stage for future chemical screens to identify new drugs with antimicrobial activity as well as other with applications in other fields such as cancer and neurodegeneration.

## Results

### Putrescine, spermidine and spermine interact with DAB to form fluorescent adducts with increasing fluorescence intensity

Previous studies by Medici et al. ^22^ and Choi et al. ^23^ have demonstrated the interaction of 1,2 diacetylbenzene (DAB) with the primary amines of molecules such as tyramine, GABA, and ethanolamine, but not those primary amines attached to α-carboxylated compounds such as serine), leading to the formation of fluorescent adducts. These fluorescent adducts could be detected using a spectrophotometer with excitation and emission spectra at wavelengths of 364 nm and 425 nm, respectively. Therefore, we examined whether a DAB-mediated fluorescence interaction with the polyamines, putrescine, spermidine and spermine could be used to measure the activity of aminopropyltransferase (APT) enzymes, which catalyze the conversion of putrescine to spermidine (SPDS activity) or spermidine to spermine (SPMS activity). First, we compared the fluorescence intensity following incubation of DAB with these polyamines as well as co-substrate, dc-SAM, or reaction product methylthioadenosine (MTA) in the absence or presence of 2 beta mercaptoethanol (β-ME) (**Fig. 1A-C**). The reaction of spermidine with 1,2 DAB/ β-ME yielded ∼3-fold higher fluorescence than that of putrescine with 1,2 DAB/ β-ME (**Fig. 1B**). On the other hand, the reaction of spermine with 1,2 DAB/ β-ME yielded ∼1.5-fold higher fluorescence than that of spermidine with 1,2 DAB/ β-ME (**Fig. 1B**). Relatively lower fluorescence intensities were detected with dc-SAM and 1,2 DAB/ β-ME, and negligible fluorescence signals were detected with MTA and 1,2 DAB/ β-ME (**Fig. 1B**). The reactivity of the mixtures of putrescine, spermidine, spermine, dc-SAM and MTA to a buffer containing DAB and lacking β-ME were also tested (**Fig. 1C**). While the overall fluorescence signals were reduced when the polyamines reacted with a detection buffer lacking β-ME, the fold difference in the fluorescence signals between putrescine-spermidine (∼3-fold) and spermidine-spermine (∼1.5-fold) remained unchanged (**Fig. 1C**).

**Figure 1.**
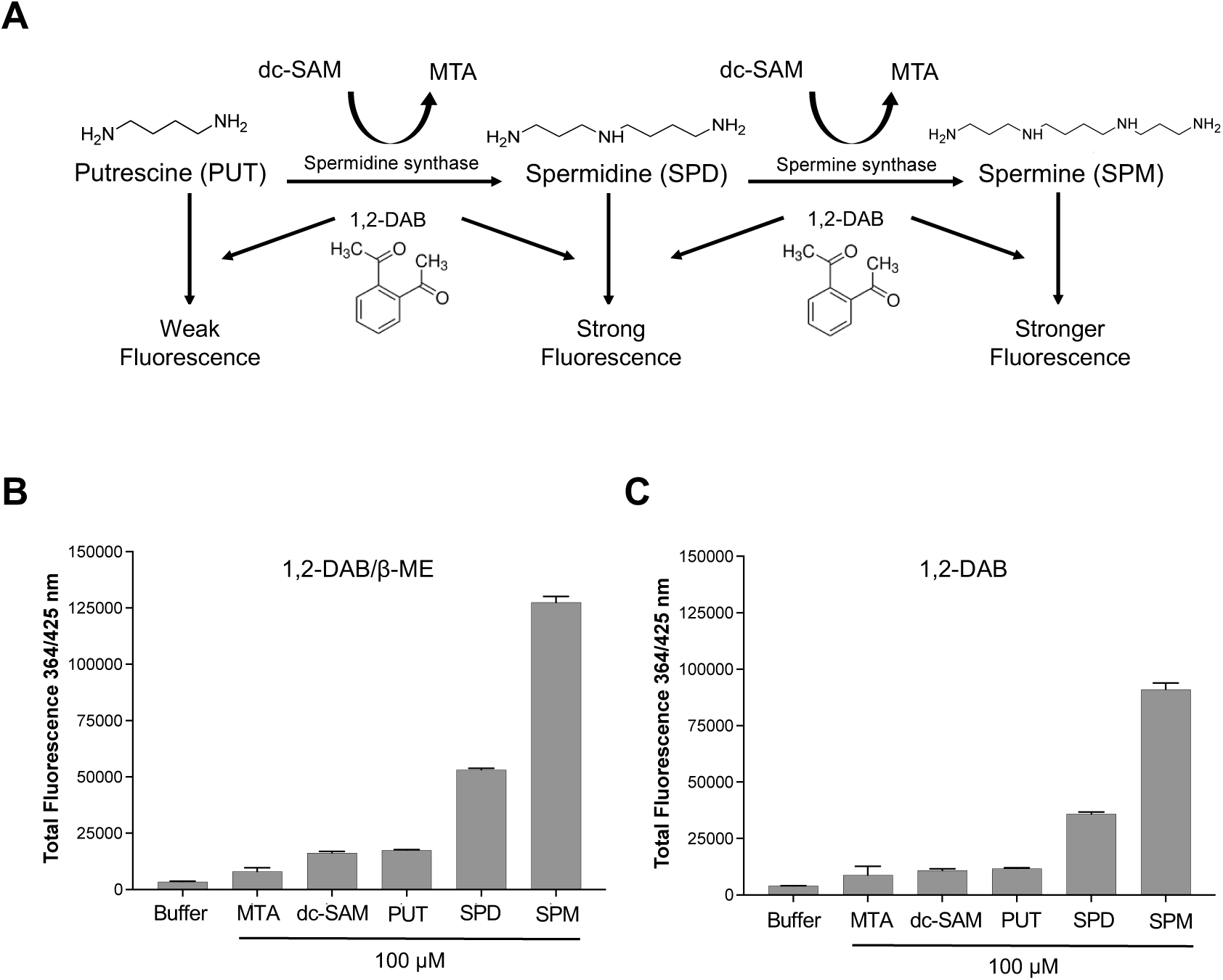
Application of 1,2 DAB/ β-ME based fluorescent assay to monitor aminopropyl transferase (APT) activity. **A**. Schematic representation of the APT reaction and the detection of the products, spermidine (SPD) or spermine (SPM), following the conversion of either putrescine (PUT) or spermidine in the presence of decarboxylated S-adenosyl methionine (dc-SAM), respectively, and subsequent interaction with 1,2 diacetyl benzene (1,2 DAB) to produce fluorescent adducts. **B-C**. Total fluorescence intensities (λ_ex_ = 364 nm and λ_em_ = 425 nm) following incubation of 1,2 DAB in the APT reaction buffer alone or supplemented with either methylthioadenosine (MTA), dc-SAM, putrescine, spermidine or spermine after 60 minutes at 22°C in the presence (**B**) or absence (**C**) of β−mercaptoethanol (β-ME). The data are presented from three independent experiments performed in triplicates, and values are mean ± S.D.

We next measured the total fluorescence (364/ 425 nm) at different concentrations (10-100μM) of putrescine, spermidine, and spermine over time (**Fig. 2A-C**). All the tested concentrations of putrescine, spermidine, and spermine showed a linear increase in the total fluorescence intensity upon reaction with 1,2-DAB/ β-ME during the first 60 minutes after which the fluorescence signal reached a plateau. At 60 min, the ratio of fluorescent intensity between spermidine and putrescine was ∼3 fold, whereas that between spermine and spermidine was ∼1.5-fold (**Fig. 2D**).

**Figure 2.**
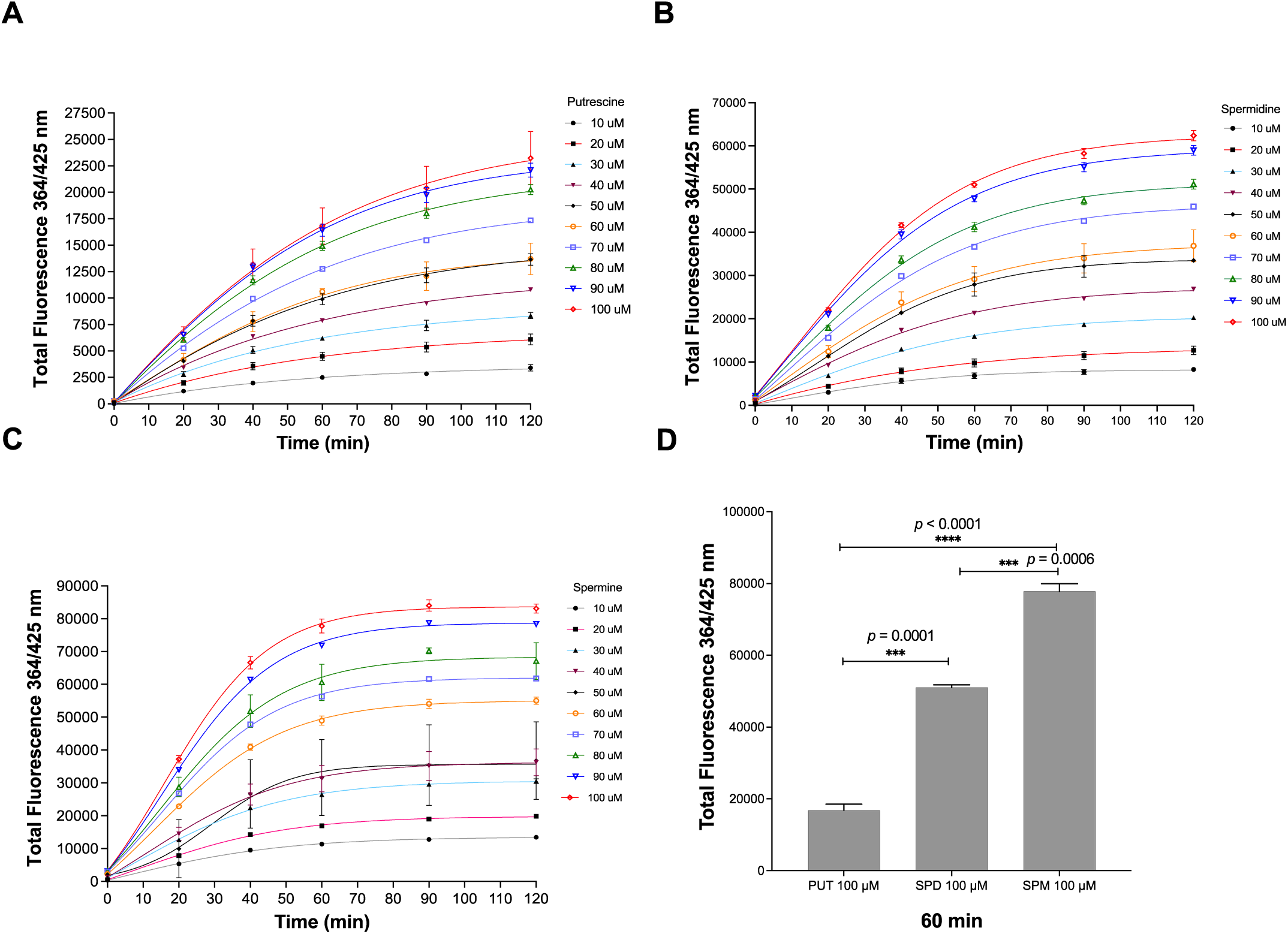
Time-dependent and concentration dependent increase in fluorescence intensity following interaction of 1,2 DAB with polyamines. **A.** Total fluorescence (λ_ex_ = 364 nm and λ_em_ = 425 nm) from 1,2 DAB/ β-ME-putrescine adducts at different concentrations of putrescine (10-100 μM) over a span of 0-120 minutes. **B**. Total fluorescence from 1,2 DAB/ β-ME-spermidine adducts at different concentrations of spermidine (10-100 μM) over a span of 0-120 minutes. **C**. Total fluorescence from 1,2 DAB/ β-ME-spermine adducts at different concentrations of spermine (10-100 μM) over a span of 0-120 minutes. **D**. Total fluorescent intensities of adducts formed between 1,2 DAB/ β-ME and 100 μM of either putrescine, spermidine, and spermine after 60 minutes of reaction. Significant difference in total fluorescent intensities were observed between PUT-SPD (p = 0.0001, PUT-SPM (p < 0.0001), and SPD-SPM (p = 0.0006). The data are presented from three independent experiments performed in triplicates, and values are mean ± S.D. The statistical significance was calculated using Welch’s t-test.

To mimic APT activities from fluorescence emission data and estimate the conversion rates in enzyme reactions, standard curves were generated using fixed ratios of the substrate and product (putrescine and spermidine or spermidine and spermine) (**Fig. S1**). The reactions were diluted in the detection buffer consisting of 1,2 DAB/ β-ME in sodium tetraborate buffer at pH 9.6, and incubated at 22°C for 60 minutes, followed by detection of fluorescence signals using spectrophotometer with excitation and emission spectra at wavelengths of 364 nm and 425 nm, respectively. The data showed direct correlation between fluorescence intensity and the rate of conversion of the substrate to the product (**Fig. S1**).

The ability to differentiate between putrescine, spermidine and spermine using 1,2 DAB, makes this assay suitable for measuring APT enzyme activities of spermidine synthases and spermine synthases.

### Activity of yeast APT enzymes, Spe3 and Spe4, using the DAB-APT fluorescence assay

To validate the use of the DAB-APT assay for measurement of APT enzyme activity, we first examined the activity of the yeast Spe3 and Spe4, which have been well-characterized at the genetic level and shown to catalyze either putrescine to spermidine or spermidine to spermine, respectively. The two enzymes were expressed in *E. coli* as N-terminal fusion proteins with the maltose binding protein (MBP) (**Fig. S2A-B**), affinity purified, and used in enzyme reactions in the presence of either putrescine or spermidine and the co-substrate dc-SAM. The reactions were stopped by the addition of the detection buffer consisting of 1,2 DAB/ β-ME in sodium tetraborate buffer at pH 9.6 and incubated at 22°C (room temperature) for 60 minutes to enable fluorescent adduct formation. Fluorescence was measured using a spectrophotometer with excitation and emission spectra at wavelengths of 364 nm and 425 nm, respectively. The yeast Spe3 catalyzes the conversion of putrescine and dc-SAM to spermidine (**Fig. 3A**). The product spermidine upon reaction with 1,2-DAB produces a strong fluorescence signal in comparison to the substrates putrescine and dc-SAM. As shown in **Figure 3B**, a time-dependent increase in fluorescence was detected when Spe3, but not Spe4, was incubated with putrescine and dc-SAM, consistent with an increase in spermidine formation (**Fig. 3B**). As a control, no fluorescence signals could be detected using heat-denatured Spe3 (Spe3_DN) or little to no increase in fluorescence could be detected when the reaction with intact Spe3 was conducted at 4°C (**Fig. 3B**). The molar amounts of spermidine formed in the Spe3 catalyzed reaction were calculated (**Fig. 3C**) from the net fluorescence data in **Fig. 3B**, using the standard curves shown in **Fig. S1B**.

**Figure 3.**
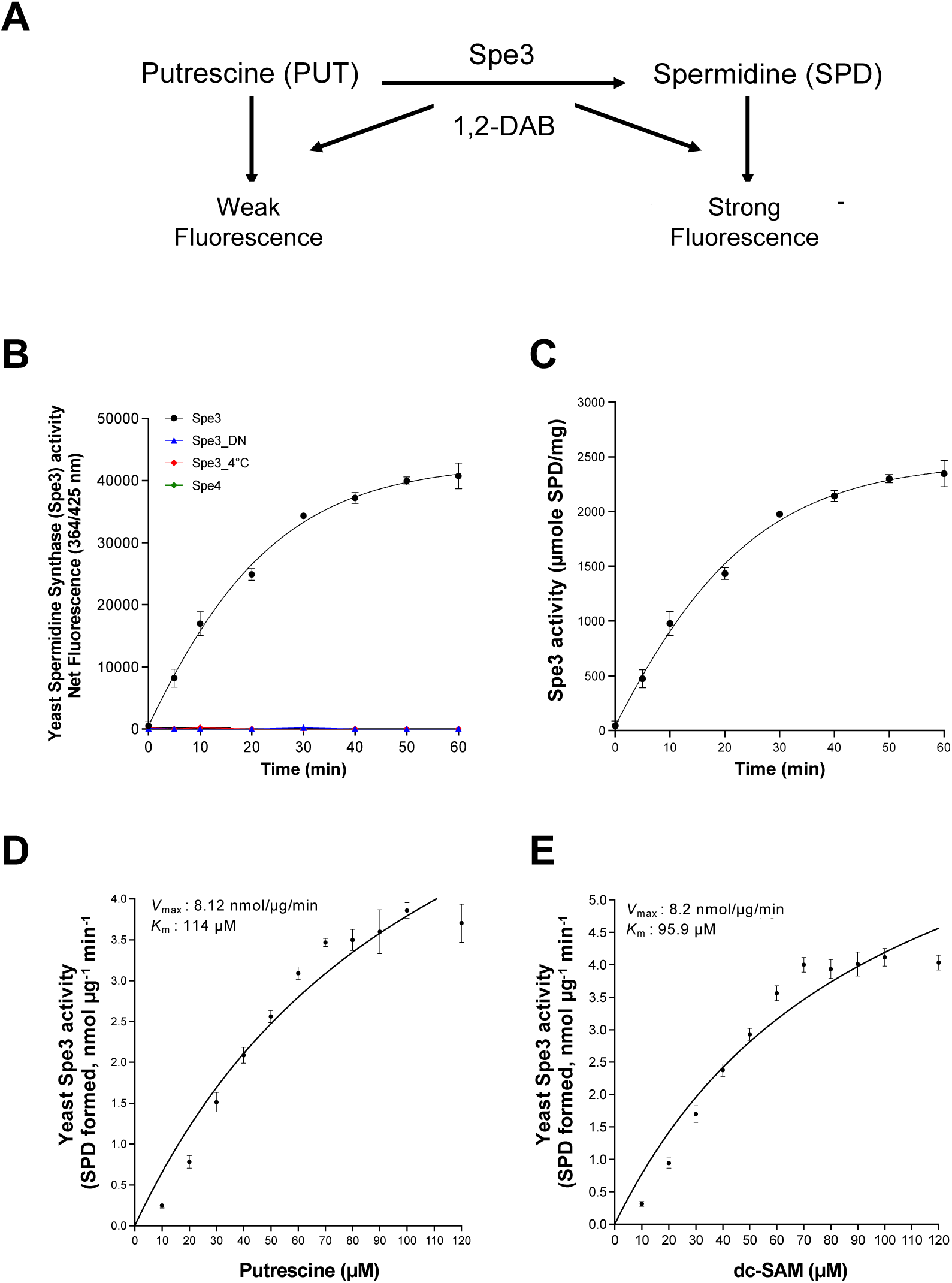
Application of the DAB-APT fluorescence assay to determine the activity and kinetics of *S. cerevisiae* spermidine synthase Spe3. **A.** Schematic representation of the enzymatic reaction catalyzed by Spe3 along with and the anticipated fluorescence signals using the DAB-APT assay. **B.** Spermidine synthase assays were conducted using affinity-purified recombinant MBP-Spe3 (20 ng/μl), heat denatured Spe3 (Spe3_DN) and 100 μM of putrescine and dc-SAM as substrate and co-substrate, respectively, at 37°C for 0-60 min. A parallel reaction at 4°C (Spe3 4°C) served as a control, the same reaction was also performed at 4°C. Purified MBP-Spe4, which lacks the ability to convert putrescine to spermidine, was included as an additional control. The spermidine synthase activity of MBP-Spe3, MBP-Spe4, and respective controls is depicted as net fluorescence intensity over time. **C.** Spermidine synthase APT activity of yeast SPE3 shown as a function of time. **D and E.** Kinetics of the Spe3 spermidine synthase activity as a function of putrescine (**D**) and dc-SAM (**E**) concentrations. Spe3-APT assays were performed with 20 ng/μl of MBP-Spe3 and varying concentrations of either putrescine or dc-SAM at 37°C for 60 minutes. *V*_max_ and *K*_m_ were determined using Michaelis-Menten kinetics in GraphPad prism. Data presented as mean ± S.D from three independent experiments, each conducted in triplicate.

This fluorescence assay for detection of APT activity provides an attractive platform for screening of novel compound libraries in a high-throughput format. Since most of the high-throughput inhibitor screens are performed using substrate concentrations near the *K*_m_ values of the substrates, we determined the kinetic constants for the yeast Spe3. For Spe3, the *K*_m_ (114 μM) and *V*_max_ (8.12 nmol/μg/min) (n=3) for putrescine as well as the *K*_m_ (95.9 μM) and *V*_max_ (8.2 nmol/μg/min) (n=3) for dc-SAM were determined (**Fig. 3D and E**) using Michaelis-Menten kinetics.

Similarly, when yeast Spe4 was used in the APT reaction with spermidine and dc-SAM as substrate and co-substrate (**Fig. 4A**), respectively, a time-dependent increase in fluorescence resulting from the formation of spermine was detected (**Fig. 4B)**. The molar amounts of spermine formed in the Spe4 catalyzed reaction **(Fig. 4C**) were calculated from the net fluorescence data in **Fig. 4B**, using the standard curves shown in **Fig. S1D**. These data demonstrate that 1,2 DAB/ β-ME is sensitive and excellent reagent for quantifying the formation of spermidine and spermine as a measure of Spe3 and Spe4 APT activity, respectively.

**Figure 4.**
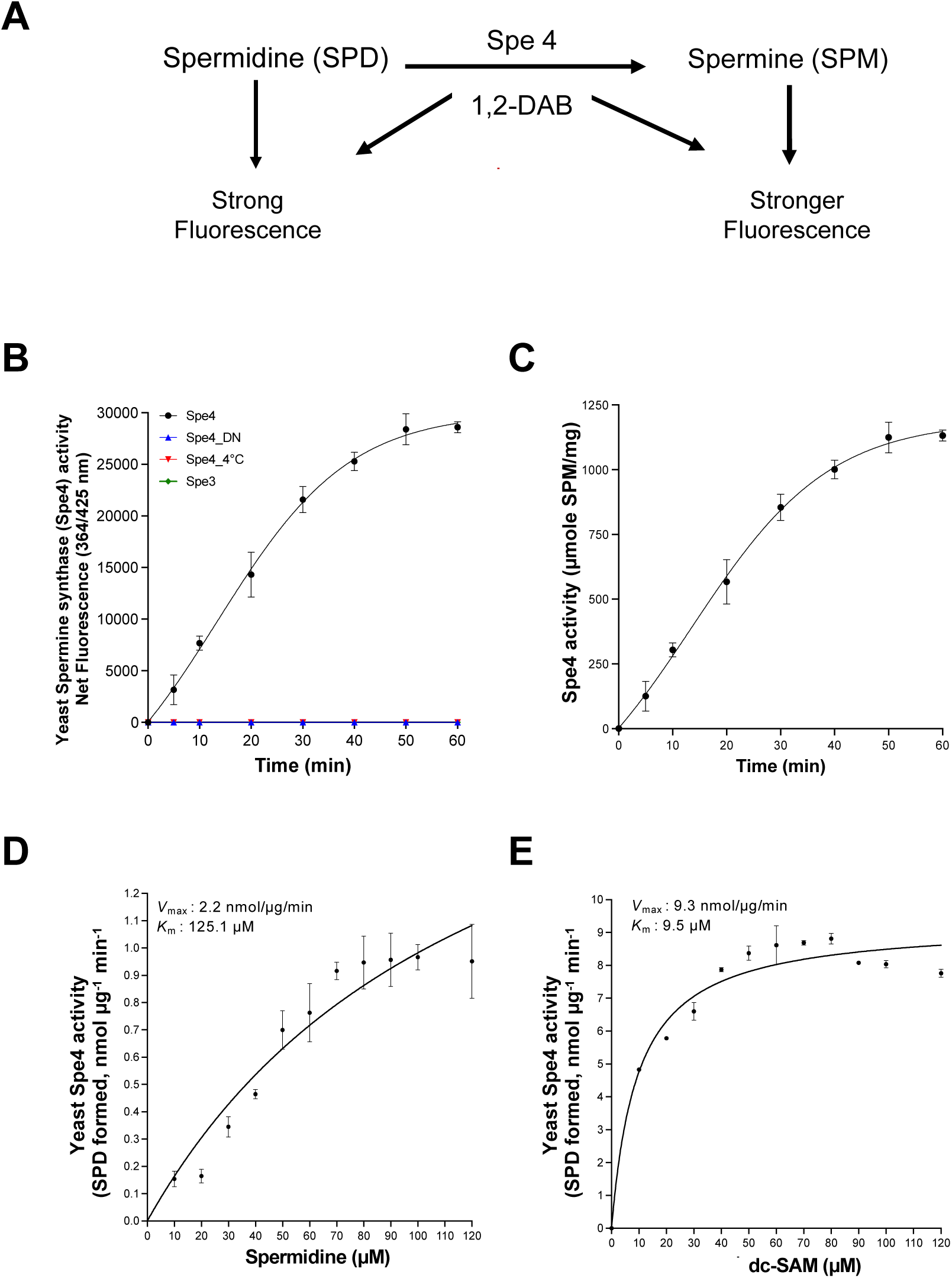
Application of the DAB-APT fluorescence assay to determine the activity and kinetics of *S. cerevisiae* spermine synthase Spe4. **A.** Schematic representation of the enzymatic reaction catalyzed by Spe4, along with and the anticipated fluorescence signals using the DAB-APT assay. **B.** Spermine synthase assays mediated by Spe4 were performed using affinity-purified recombinant MBP-Spe4 (20 ng/μl), heat-denatured SPE4 (Spe4_DN), along with 100 μM of spermidine and dc-SAM as substrates at 37°C for 0-60 minutes. A control reaction was conducted at 4°C (Spe 4°C). Spe3, which lack the ability to convert spermidine to spermine was used as an additional control. The spermine synthase activity of MBP-Spe4, MBP-Spe3, and controls is depicted as net fluorescence intensity over time. **C.** Spermine synthase activity of yeast Spe4 represented as molar amounts of spermidine formed over time. **D and E.** Kinetics of the Spe4 spermine synthase activity as a function of spermidine (**D**) and dc-SAM (**E**) concentrations. Spe4-APT assays were performed with 20 ng/μl of MBP-Spe3 and varying concentrations of either spermidine or dc-SAM at 37°C for 60 minutes. *V*_max_ and *K*_m_ were determined using Michaelis-Menten kinetics in GraphPad prism. Data presented as mean ± S.D from three independent experiments, each conducted in triplicate.

The kinetics for Spe4 were also performed, where the *K*_m_ (125.1 μM) and *V*_max_ (2.2 nmol/μg/min) (n=3) for spermidine as well as the *K*_m_ (9.5 μM) and *V*_max_ (9.3 nmol/μg/min) (n=3) for dc-SAM were determined (**Fig. 4D and 4E**).

### The 1,2-DAB/ β-ME–based fluorescence assay can measure the activity of the purified *P. falciparum* SPDS enzyme

We further used the 1,2-DAB/ β-ME-APT assay, to determine the APT activity and kinetic properties the *P. falciparum* spermidine synthase, PfSPDS, using a recombination MBP-PfSPDS enzyme (**Fig. 5**). The enzyme was previously examined for its catalytic activity using ^14^C-putrescine ^24^, and shares high sequence similarity with APT enzymes from *E. coli*, *S. cerevisiae*, and *H. sapiens*, especially in the catalytic site with the key residues aspartate 127, glutamate 147 and aspartate 196 in PfSPDS all found to be conserved among other well-known APT enzymes (**Fig. 5A and** S4). Using the DAB-APT assay, PfSPDS was found to catalyze the conversion of putrescine to spermidine as determined by the time-dependent increase in fluorescence following incubation with DAB (**Fig. 5B**). As a control, heat-inactivated PfSPDS failed to convert putrescine to spermidine (**Fig. 5B**). To further validate these results, the key conserved residues, D127, E147 and D196, required for APT activity in PfSPDS were mutated to alanine and the activity of the mutant PfSPDS enzymes was tested (**Fig. 5A-B**, **Fig. S2C-F**). All three mutated enzymes, PfSPDS^D127A^, PfSPDS^E147A^ and PfSPDS^D127A,E147A, D196A^, failed to convert putrescine to spermidine (**Fig 5B**). The molar amounts of spermidine formed in the PfSPDS catalyzed reaction (**Fig. 5C**) were calculated from the net fluorescence data in **Fig. 5B**. The assay was subsequently used to determine the kinetic parameters of the PfSPDS enzyme. The calculated *K*_m_ and *V*_max_ (n=3) for dc-SAM were 51.89 μM, and 8.336 nmol/μg/min (**Fig. 5D**), and the *K*_m_ and *V*_max_ (n=3) for putrescine were 66.49 μM, and 9.002 nmol/μg/min (**Fig. 5E**). Conversely, the DAB-APT assay demonstrated that PfSPDS lacks a spermine synthase activity (**Fig. S5**), suggesting that, similar to other eukaryotes, the spermine synthase activity in *P. falciparum* is most likely catalyzed by a separate enzyme.

**Figure 5.**
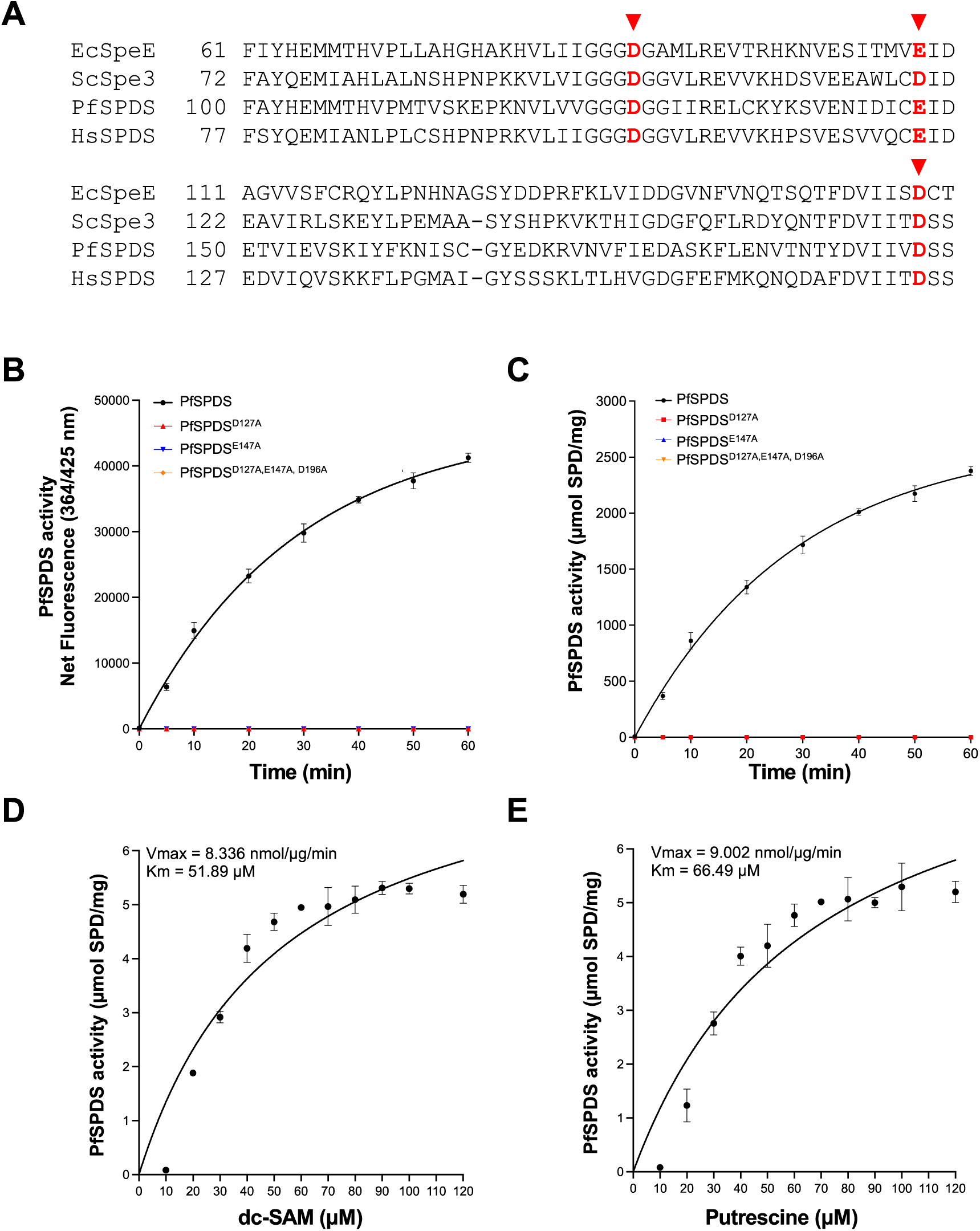
Application of the DAB-APT fluorescence assay to determine the activity and kinetics of the *P. falciparum* PfSPDS enzyme. **A.** Sequence alignment of PfSPDS with *E. coli*, *S. cerevisiae*, and *H. sapiens* spermidine synthases. Key amino acids critical for enzyme activity are highlighted in red. **B.** Spermidine synthase reactions were conducted using affinity-purified recombinant MBP-PfSPDS (20 ng/μl), heat-denatured PfSPDS (PfSPDS_DN), and 100 μM of putrescine and dc-SAM as substrate and co-substrate, respectively, at 37°C for 0-60 minutes. A parallel reaction with active PfSPDS at 4°C (PfSPDS 4°C) served as a control. PfSPDS mutants, PfSPDS^D127A^, PfSPDS^E147A^, and PfSPDS^D127A, E147A, D196A^ were used to validate the importance of these residues in PfSPDS catalytic activity. The spermidine synthase activity of MBP-PfSPDS, PfSPDS mutants, and controls is depicted as net fluorescence intensity over time. **C.** Spermidine synthase activity of PfSPDS shown as a function of time. **D and E.** Kinetics of PfSPDS spermidine synthase activity as a function of dc-SAM (**D**) or putrescine (**E**) concentrations. PfSPDS-APT assays were conducted with PfSPDS and varying concentrations of either dc-SAM or putrescine at 37°C for 60 minutes. *V_max_* and *K_m_* values were determined using Michaelis-Menten kinetics in GraphPad Prism. Data are presented as mean ± S.D. from three independent experiments, each conducted in triplicate.

### The DAB-APT assay is amenable to HTS screening

The adaptability of 1,2 DAB/ β-ME-APT assay to a 96-well plate format for conducting inhibitor screening in HTS platforms was evaluated using trans-4-methylcyclohexylamine (4MCHA), a known potent inhibitor of spermidine synthase. 4MCHA has been demonstrated to compete with putrescine binding sites on spermidine synthase enzymes^25^. The putrescine to spermidine APT activity of *S. cerevisiae* Spe3 and PfSPDS was determined in 96-well format in a 100 µl volume in the absence or presence of increasing concentrations of 4MCHA (0-100 μM) and spermidine formation was determined by measuring fluorescence intensity following addition of DAB (**Fig. S3**). For Spe3 and PfSPDS enzymes, the calculated EC_50_ values of 4MCHA were ∼ 7.8 μM and ∼ 6 μM, respectively (**Fig. 6A**). The inhibition constants for the compound were determined to be *K_i_* = 0.82 μM for Spe 3 and *K_i_* = 0.64 μM for PfSPDS enzymes (**Fig. 6B-C**). The kinetic parameters *K*_m_ and *V*_max_ of the inhibition curves obtained with different concentration of 4MCHA indicates a competitive inhibition by 4MCHA, which supports the previous findings where 4MCHA has been demonstrated to compete with putrescine for binding to the enzyme ^25^.

**Figure 6.**
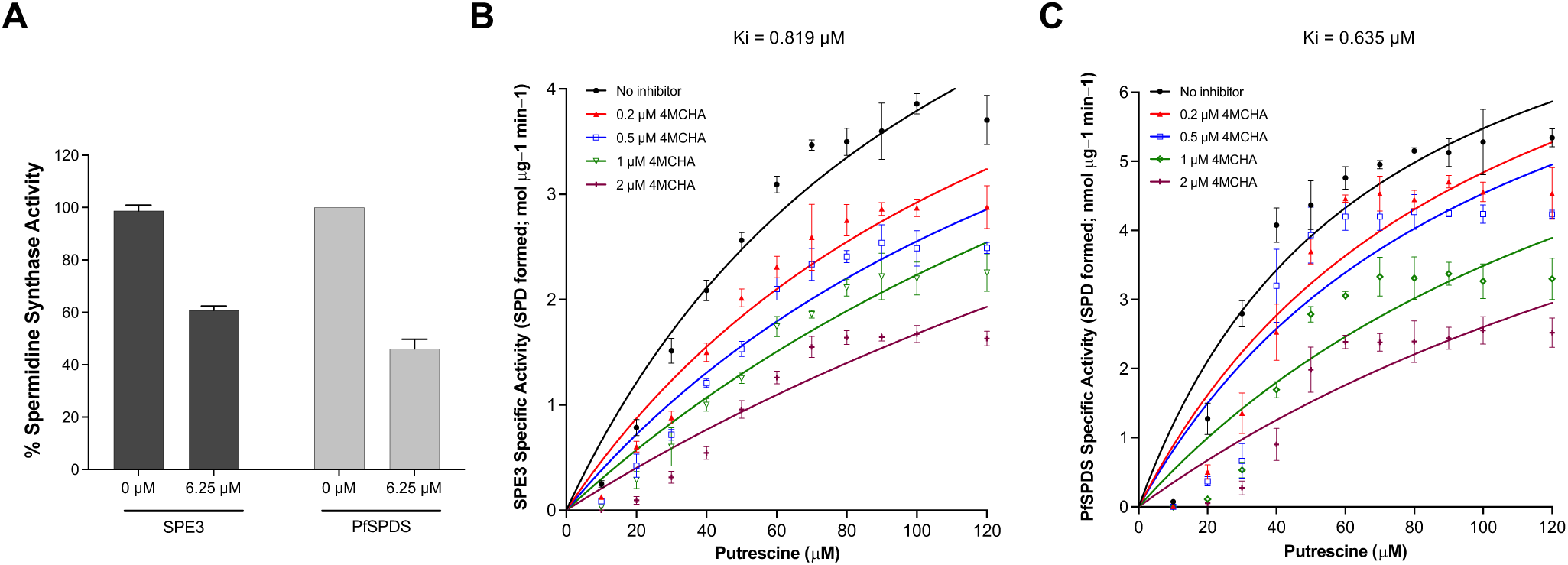
Inhibition of *S. cerevisiae* Spe3 and *P. falciparum* PfSPDS activity by 4MCHA. **A.** Total fluorescence intensity as a function of Spe3 and PfSPDS activity in the absence or presence of 4MCHA (6.25 µM (its calculated EC_50_)). **B** and **C**. Determination of inhibition constants (Ki) values for inhibition of *S. cerevisiae* Spe3 (**B**) and *P. falciparum* PfSPDS (**C**) spermidine synthase activity by 4MCHA. All data are presented as mean ± S.D from three independent experiments, each conducted in triplicate.

Amenability of the DAB-APT assay to HTS was determined for the yeast Spe3 and Spe4 enzymes by calculating the signal to background (S/B) and *Z*’ score as detailed in the Methods. From these analyses, the yeast Spe3 had S/B of 773.6 and a *Z*’ score of 0.83, and the yeast Spe4 had S/B ratio of 242.2 and a *Z*’ score of 0.88. These data validate the assay’s suitability for high-throughput screening and may be used to search for novel inhibitors in available chemical libraries for various indications.

## Discussion

In this study, we describe a novel fluorescence assay utilizing 1,2-diacetylbenzene (DAB) and β-mercaptoethanol (β-ME) for measurement of the activity of aminopropyltransferases (APTs). The assay builds upon previous successful use of DAB to measure the activity of phosphatidylserine decarboxylase enzymes that convert phosphatidylserine to phosphatidylethanolamine due to interaction of DAB with ethanolamine but not serine ^23^. Mass spectrometry analysis demonstrated the formation of an iso-indole oxathiolane adduct responsible for the fluorescence signal ^23^. Our studies demonstrated that DAB interacts with primary amines of the polyamines, putrescine, spermidine and spermine, to form fluorescent adducts, likely through a similar chemical process as with ethanolamine ^23^. Although all three polyamines carry two primary amine groups that are available for reaction with 1,2 DAB, our data showed differential fluorescence signals for putrescine, spermidine, and spermine (**Fig. 1B)**. We hypothesize that the higher fluorescence signals observed with spermine (tetramine) versus spermidine (triamine) and putrescine (diamine) are due to increased polyamine chain length that separates the two primary amine groups in these molecules. The adducts formed between primary amine groups of putrescine and 1,2 DAB are thus in close proximity and potentially quench the fluorescence signals due to steric hindrance. As the polyamine chain length increases, the primary amine groups become far from each other leading to higher fluorescence intensity.

Our data also showed that the addition of β-ME enhanced the sensitivity of the assay, enabling the detection of subtle changes in APT enzyme activity. This suggests a potential mechanism involving β-ME in facilitating the interaction between DAB and polyamines, thereby amplifying the fluorescence signal. Our findings highlight the importance of optimizing assay conditions to maximize sensitivity and specificity, essential for accurate enzyme activity measurements.

To validate the DAB-based assay for measuring the activity of APT enzymes, we investigated the activity of the *S. cerevisiae*, Spe3 and Spe4, enzymes as well as the spermidine synthase (SPDS) from *P. falciparum*. The assay demonstrated high sensitivity and specificity, enabling the quantification of spermidine and spermine formation over time. Moreover, determination of the kinetic parameters of these enzymes provided insights into their catalytic properties, essential for understanding polyamine metabolism and enzyme function.

Importantly, our assay exhibited suitability for inhibitor screening in high-throughput platforms. By utilizing known inhibitors, such as trans-4-methylcyclohexylamine (4MCHA), we successfully determined their inhibitory potency against APT enzymes. The competitive inhibition observed with 4MCHA underscores the potential of our assay in identifying novel inhibitors targeting polyamine metabolism, with implications for therapeutic intervention in diseases such as cancer and parasitic infections.

In conclusion, our study presents a robust and versatile fluorescence assay for accurately measuring APT enzyme activity. The assay’s sensitivity, specificity, and suitability for high-throughput screening make it a valuable tool for drug discovery and polyamine metabolism research. We anticipate that our findings will pave the way for future studies exploring the role of APT enzymes in health and disease, as well as the development of novel therapeutics targeting polyamine metabolism pathways.

## Methods Materials

*S. cerevisiae SPE3* and *SPE4*, and *P. falciparum* SPDS were codon-optimized for expression in *E. coli*, chemically synthesized and cloned into pMAL-c4x-1-H(RBS) plasmid by GenScript. Putrescine (P5780-5G), spermidine (S0266-1G), spermine (S4264-1G), 5-Deoxy-5-methylthioadenosine (260585), 1,2-Diacetylbenzene (DAB) (242039) were purchased from Millipore Sigma. Decarboxylated S-adenosyl methionine was purchased from BOC Sciences, USA, and 2-Mercaptoethanol (1610710) was purchased from Bio-Rad, USA.

### Expression and purification of MBP-tagged SPE3, SPE4 and PfSPDS

The *SPE3*-pMAL-c4x-1-H(RBS), *SPE4*-pMAL-c4x-1-H(RBS) and *PfSPDS*-pMAL-c4x-1-H(RBS) plasmids were transformed into *Rosetta (DE3) E. coli* cells ((Fisher Scientific, 713973). Two clones from each were selected and tested for expression of MBP-tagged Spe3, Spe4, and PfSPDS. Briefly, each clone was inoculated into 5 mL of Luria broth ^24^ containing ampicillin (50 μg/ml) and allowed to grow overnight at 37°C incubator shaker (200 rpm). The following day, secondary cultures were initiated using the primary cultures and grown to 0.6 OD_600_. Following this, 0.5 mM isopropyl-thiogalactopyranoside (IPTG) was added and the cultures were shifted from 37°C to 16°C and allowed to grow for 16 hours. The cultures were collected by centrifugation at 5000 g for 5 min. The pellets were resuspended in 1X Laemmli sample buffer (1610737EDU; Bio-Rad), boiled at 95°C for 10 min and centrifuged at 10,000 g for 5 min. The supernatants from uninduced and induced samples were run on 4-20% SDS-PAGE (4561096; Bio-Rad) to check protein expression using Coomassie staining.

For the purification of recombinant MBP-tagged Spe3, MBP-Spe4, MBP-PfSPDS, 500 mL culture for each protein was grown in LB medium containing ampicillin (50 μg/ml) and induced with 0.5 mM IPTG. The bacterial pellets were harvested ∼12 hours after induction and resuspended in lysis buffer (25 mM Tris-HCl pH 8.0, 500mM NaCl, 0.5% glycerol, and 50 mM L-arginine, DNase 250 UL/μl, protease inhibitor cocktail, 0.002% 3-((3-cholamidopropyl) dimethylammonuium-1-propane sulfonate (CHAPS) and disrupted by sonication on ice (Omni Sonic Ruptor 400 Ultrasonic Homogenizer) by 15 sec burst at 70% amplitude, 5 times, with 30 sec cooling intervals. The bacterial lysates were centrifuged at 16,000 x g for 20 min and the supernatants containing the recombinant MBP-Spe3, MBP-Spe4, and MBP-PfSPDS were collected. Next, three columns containing 500 μL of amylose magnetic beads (NEB, E8021L) each were packed and equilibrated with column buffer (200 mM NaCl, 20 mM Tris-HCl, 1 mM EDTA, 1 mM DTT). The supernatants were incubated with the amylose resin for 2.5 hour at 4°C with end-to-end shaking. Following this, the columns were washed with 10 volumes of the column buffer and the protein was eluted using 10 mM maltose (M75-100; Fisher Scientific). The purified recombinant MBP-Spe3, MBP-Spe4, and MBP-PfSPDS were run on SDS-PAGE and stained with Coomassie blue to check the purity. The protein concentration was determined using nanodrop (Biotek SynergyMX with take3 plate; TAKE3-SN).

### Fluorescence based APT assay

The APT enzyme reaction and subsequent detection of the products (spermidine and spermine) using 1,2 DAB/β-ME was performed in the following sequential steps. In the first step, recombinant Spe3, Spe4, and PfSPDS were used in the aminopropyl transferase activity assay. For determining the spermidine synthase activity of recombinant Spe3 and PfSPDS, the enzyme reactions were set up containing 0.1 mM of dc-SAM (BOC Biosciences), 0.1 mM putrescine, 1mM EDTA, 1mM DTT, 1 μg BSA, 1μg of either Spe3 or PfSPDS, and 50 mM potassium phosphate buffer (pH 7.5), in 50 μl total volume. The enzyme reaction was incubated at 37°C for 60 minutes. For determination of spermine synthase activity of Spe4, the enzyme reactions were set up containing 0.1 mM of dc-SAM, 0.1 mM spermidine, 1mM EDTA, 1mM DTT, 1 μg BSA, 1μg Spe4, and 50 mM potassium phosphate buffer (pH 7.5), in 50 μl total volume. The enzyme reaction was incubated at 37°C for 60 minutes. Subsequently, 35 μL of the above enzyme reactions were mixed with 85 μL detection buffer (1.75 mM β-mercaptoethanol, 64.5 mM sodium tetraborate buffer (pH 9.6), 0.22 mM potassium phosphate buffer, 1.48 mM 1,2 diacetyl benzene) in a 96-well black clear bottom plate (265301, ThermoFisher Scientific) and allowed to incubate at 22°C (room temperature) for 60 minutes. Following this, total fluorescence intensities (λ_ex_ = 364 nm and λ_em_ = 425 nm) were measured using a plate reader (Biotek Synergy H1, Agilent) and data was analyzed in GraphPad prism.

### Data analysis to determine *Z*’ score

The signal to background (S/B) ratios for the yeast Spe3 and Spe4 were calculated using the following equation:

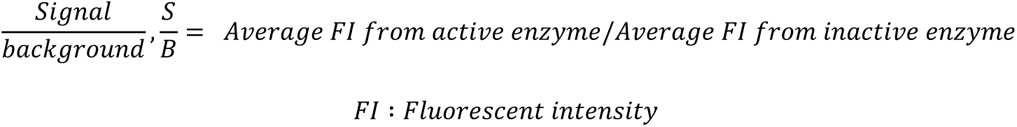

The coefficient of variation (CV) were calculated using the following formula:

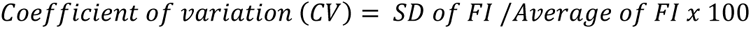

The *Z*’ factor was calculated using the following equation:

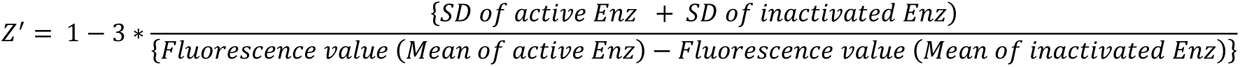

## Data availability

All data are contained in the article and the supplemental data files.

### Author contributions

P.S., J.Y.C., C.B.M. conceptualization; P.S., J.Y.C. data curation; P.S., J.Y.C., C.B.M. writing-review, and editing; C.B.M. funding acquisition.

### Funding and additional information

CBM research is supported by NIH grants AI138139, AI152220, AI123321 and AI136118, and AI153100, and the Steven and Alexandra Cohen Foundation [Lyme 62 2020 to CBM].

### Conflict of interest

The authors no conflict of interest, financial or otherwise.

---

**Figure S1.**
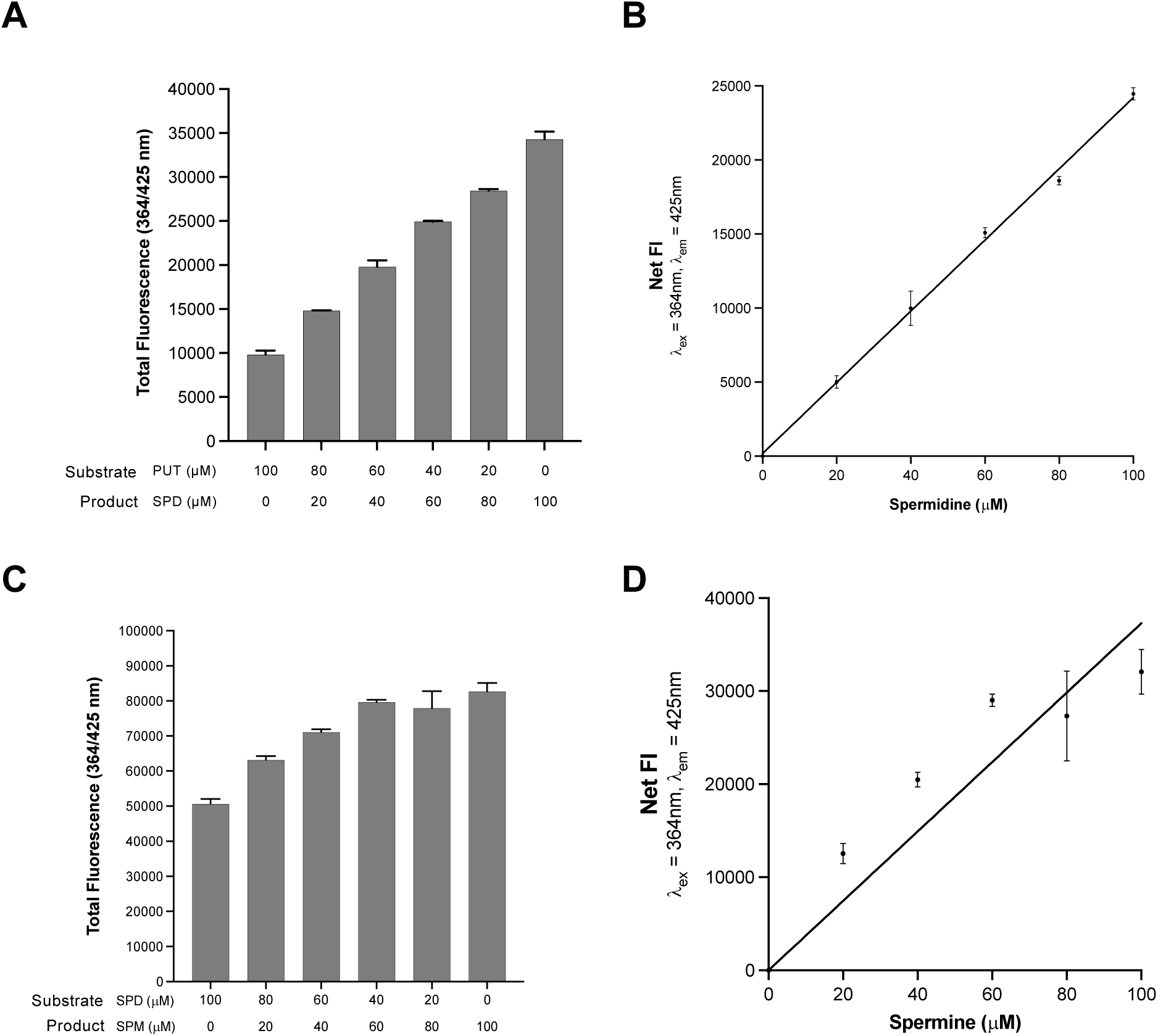
Changes in the ratios of putrescine-spermidine and spermidine-spermine produce quantitative changes in 1,2-DAB/β-ME fluorescence. **A**. Putrescine and spermidine were mixed in different ratios as indicated in the figure (decreasing concentration of putrescine and increasing concentrations of spermidine), and the total PUT+SPD concentration was maintained at 100 μM. The total fluorescence intensity was measured at 364 nm (excitation)/ 425 nm (emission) after 1 h of incubation with 1,2-DAB/β-ME. **B.** The net fluorescent intensity (FI) was calculated from data in figure S1A. A linear increase in net fluorescence intensity of spermidine with increasing concentration was observed. **C.** Spermidine and spermine were mixed in different ratios as indicated in the figure (decreasing concentration of spermidine and increasing concentrations of spermine), and the total SPD+SPM concentration was maintained at 100 μM. The total fluorescence intensity was measured at 364 nm (excitation)/ 425 nm (emission) after 1 h of incubation with 1,2-DAB/β-ME. **D.** The net fluorescent intensity (FI) was calculated from data in figure S1C. A linear increase in net fluorescence intensity of spermine with increasing concentration was observed. The data are from three independent experiments conducted in triplicates, with error bars denoting mean ± S.E.

**Figure S2.**
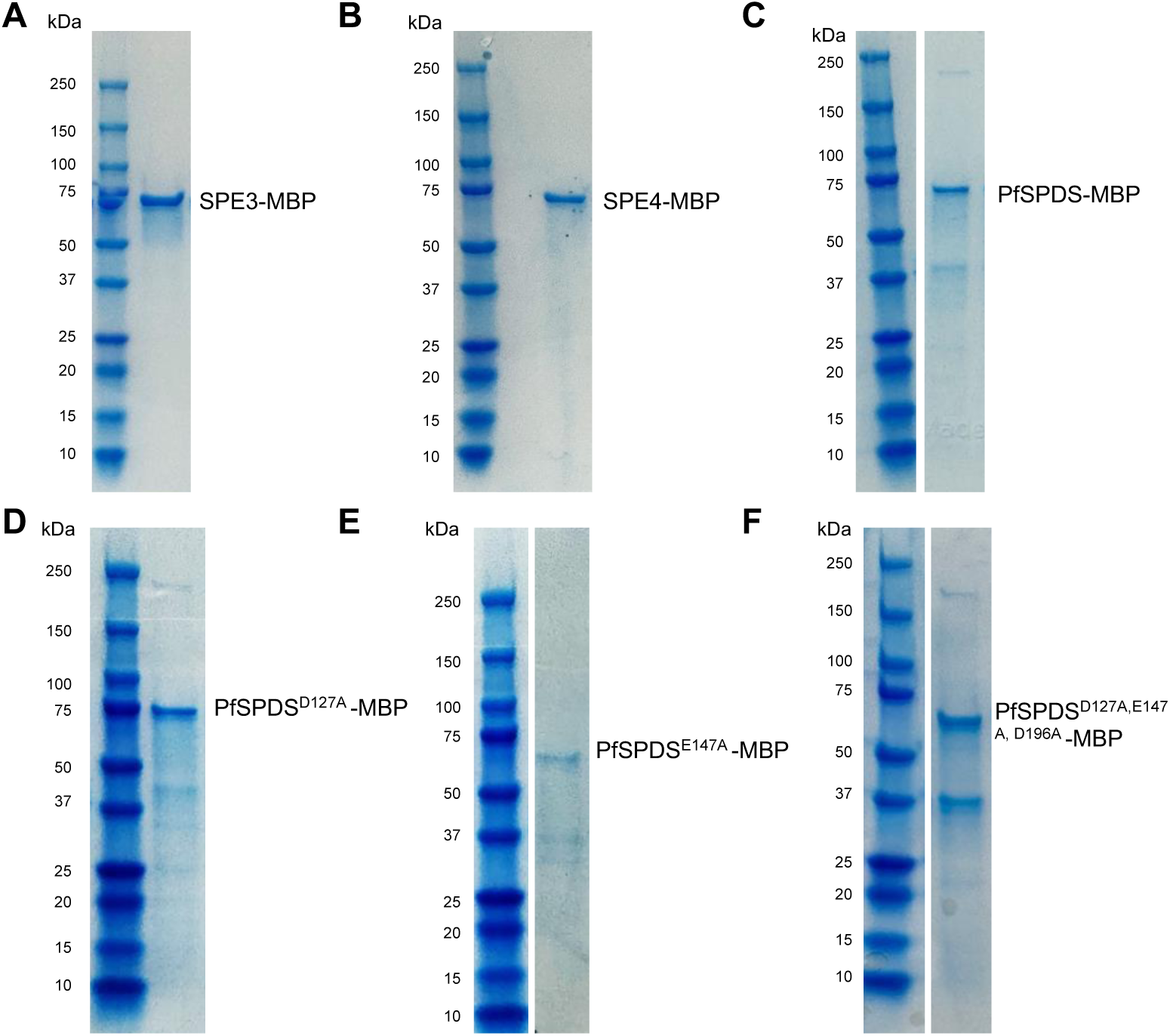
Purification of recombinant MBP-tagged SPE3, SPE4, PfSPDS, PfSPDS^D127A^, PfSPDS_E147A_, and PfSPDS^D127A, E147A,D196A^.

**Figure S3.**
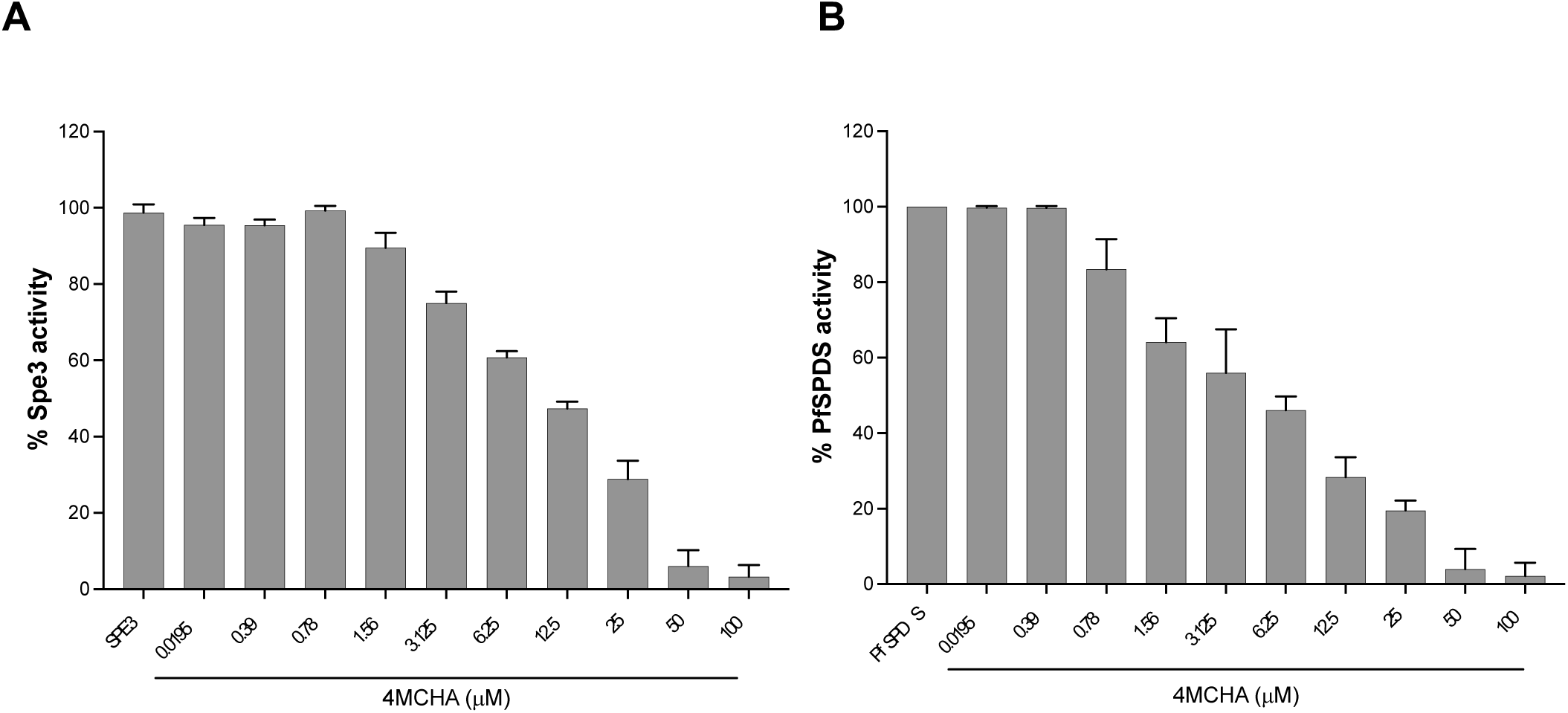
Dose dependent inhibition of yeast Spe3 and PfSPDS activity with trans-4-methylcyclohexylamine (4MCHA). **A.** A dose-dependent decrease in the activity of SPE3 with increasing concentrations of 4MCHA (0.0195μM-100 μM). **B.** A dose-dependent decrease in the activity of PfSPDS with increasing concentrations of 4MCHA (0.0195μM-100 μM). The data are from three independent experiments conducted in triplicates, with error bars denoting mean ± S.E.

**Figure S4.**
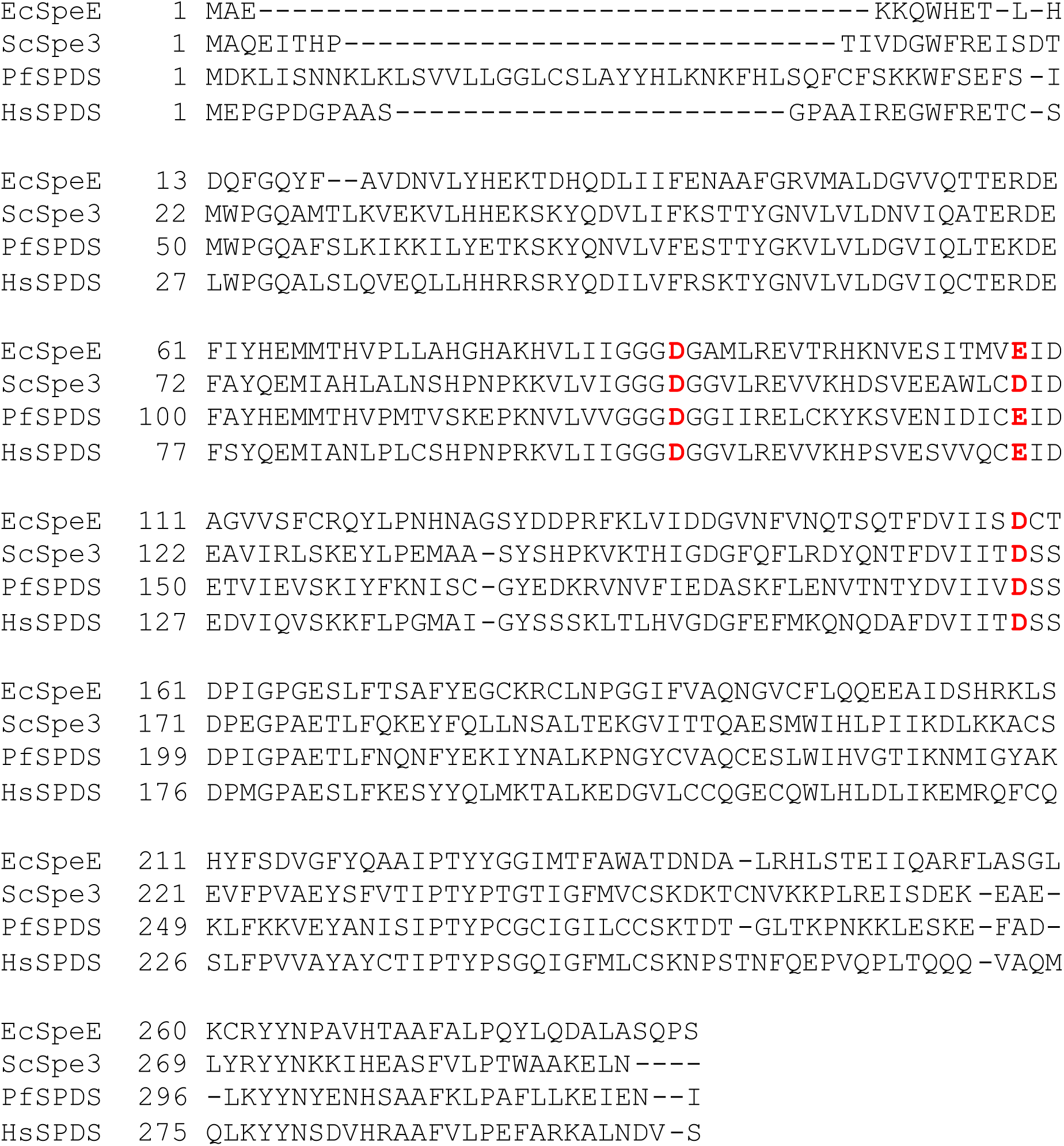
Multiple sequence alignment of *P. falciparum* spermidine synthase (PfSPDS) with spermidine synthase enzymes from *E. coli, S. cerevisiae*, and *H. sapiens*.

**Figure S5.**
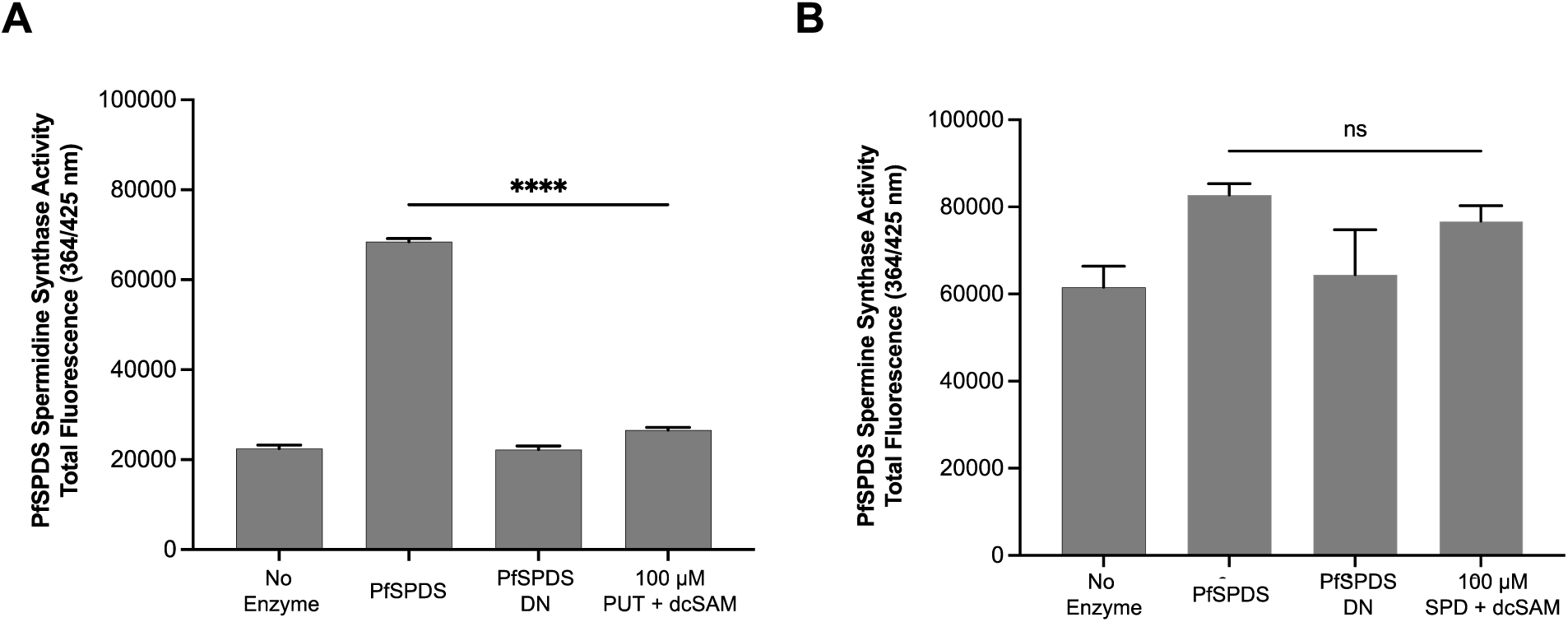
PfSPDS possesses spermidine synthase activity but lacks spermine synthase activity. **A.** Recombinant MBP-tagged PfSPDS was used in 1,2-DAB-APT assay to test the spermidine synthase activity, using putrescine (100 μM) and dcSAM (100 μM) as substrates. The enzyme reactions containing no enzyme or heat denatured enzyme (PfSPDS DN) were used as controls. A mock reaction containing 100μM putrescine and dc-SAM was also used as a control. The total fluorescence (364nm/425nm) from the above reactions were measured. A significant increase in the total fluorescence intensity was observed in the PfSPDS catalyzed reaction in comparison to mock reaction containing only substrates. The data are from three independent experiments conducted in triplicates, with error bars denoting mean ± S.E. **B**. Recombinant MBP-tagged PfSPDS was used in 1,2-DAB-APT assay to test the spermine synthase activity, using spermidine (100 μM) and dcSAM (100 μM) as substrates. The enzyme reactions containing no enzyme or heat denatured enzyme (PfSPDS DN) were used as controls. A mock reaction containing 100μM of putrescine and dc-SAM was also used as a control. No significant difference in the total fluorescence intensity was observed in the PfSPDS catalyzed reaction in comparison to mock reaction containing only substrates. The data are from three independent experiments conducted in triplicates, with error bars denoting mean ± S.E.

## Notes

### Competing Interest Statement

The authors have declared no competing interest.

